# Identification of essential regulatory elements in the human genome

**DOI:** 10.1101/444562

**Authors:** Alex Wells, David Heckerman, Ali Torkamani, Li Yin, Bing Ren, Amalio Telenti, Julia di Iulio

## Abstract

The identification of essential regulatory elements is central to the understanding of the consequences of genetic variation. Here we use novel genomic data and machine learning techniques to map essential regulatory elements and to guide functional validation. We train an XGBoost model using 38 functional and structural features, including genome essentiality metrics, 3D genome organization and enhancer reporter STARR-seq data to differentiate between pathogenic and control non-coding genetic variants. We validate the accuracy of prediction by using data from tiling-deletion-based and CRISPR interference screens of activity of *cis*-regulatory elements. In neurodevelopmental disorders, the model (*ncER*, non-coding Essential Regulation) maps essential genomic segments within deletions and rearranged topologically associated domains linked to human disease. We show that the approach successfully identifies essential regulatory elements in the human genome.

---

There is rapid improvement in the understanding of the human genome, the organization of function, and the consequences of human genetic variation. This understanding enables multiple innovations in medical genetics. For example, exome sequencing accelerates the diagnosis and contributes to the clinical management of rare genetic disorders. However, exome sequencing only examines less than 2% of the genome sequence and the current diagnostic yield stands at around 30%^1-4^. The role of genetics in the remaining 70% of rare disorders is at present unknown, and the exact mechanisms of disease remain largely unexplored. One potential mechanism is through perturbation of important regulatory regions of the genome. Recent efforts at extending the search space from the coding to the immediate regulatory regions show that new pathogenic variants can be identified in a small fraction of cases^5,6^. In parallel, there are recent examples of diseases that implicate distal enhancers and changes in the 3D genome structure^6,7^. Thus, the next milestones in the interpretation of the human genome sequence will emerge from the analysis of functional consequences of genetic variants in the non-protein coding (here in referred to as non-coding) genome – the remaining 98% of genome sequence that includes the regulatory machinery. As previously done for coding genes^8^, we define a regulatory element as essential when loss of its function may compromise viability of the individual or results in profound loss of fitness or in disease.

Interpretation of the non-coding genome requires the identification of landmarks, features and structures – analogous to the same first principles that aid the interpretation of the coding genome. This translates to the characterization of regulatory genomic elements, rules of functional essentiality and redundancy, and the interpretation of the organization of the genome. Genome-wide epigenomic maps have revealed hundreds of thousands regions showing signatures of enhancers, promoters, and other gene-regulatory elements^9^. However, the high-resolution dissection of driver nucleotides as well as the characteristics of essentiality of those regions remain limited at present^10^. There are multiple sources of data (biochemical, genetic, evolutionary) that convey functional information on the non-coding genome^11^. These data are used by different scoring algorithms ^12-21^ that aim at ranking variants according to their predicted deleteriousness. The accuracy of these methods typically increases through collective integration of multiple models, eg. ensemble-based classifiers_22_. In addition, there is an opportunity to increase the precision of functional and deleteriousness prediction by learning from novel data sources – in particular, resources that have not been included in previous analyses. Those novel sources of data include studies of the patterns of human-specific constraints that are revealed by population genomic analyses^23^, analyses of 3D organization of the genome (eg., promoter capture Hi-C)^24,25^, and from high throughput screens of enhancer function^26^. In this work we implement state-of-the art machine learning tools to rank-classify the most essential regulatory elements of the non-coding genome with an emphasis on the contribution of new data modalities. We then use tiling array deletion and CRISPR interference (CRISPRi) data to assess the accuracy and functional relevance of the predictions. Lastly, we use the new predictive tools to identify essential regulatory regions associated with human disease – in structural variants associated with the autism spectrum disorder (ASD) and in developmental disorders resulting from the disruption of topological associated domains (TADs). The study design is summarized in **Suppl. Figure S1**.

## Training a model to identify essential regulatory elements

To train a supervised machine learning model, we included non-coding pathogenic variants from ClinVar^27^ and Human Gene Mutation Database (HGMD)^28^ (N= 1,095, **Methods**). The set of control variants was built by using all variants from gnomAD (http://gnomad.broadinstitute.org/) with allele frequencies greater than 1% across populations and sub-selecting (N=8,093) those that matched the pathogenic variant set based on distance to splice sites and genomic element distribution (**Methods**). For validation, we used non-coding pathogenic variants not included in the original dataset: an independent set of manually curated non-coding Mendelian pathogenic variants (n=425)^23^, and a new release of ClinVar and HGMD (total of N=599, including N=245 mapping to the ncRNAs); **Methods**.

We trained an XGBoost model (https://xgboost.readthedocs.io), an implementation of gradient-boosted decision trees consisting of a collection of decision trees, where a node in a single decision tree splits the training data into subsets (deleterious versus benign). During testing, new variants with the same feature sets were given to each tree to make a prediction (essential or non-essential). The outputs of each tree were combined (“ensembling”) to generate a final prediction. Each variant in the dataset was annotated with 38 features from 4 major categories (**Suppl. Table S1, Methods**). (i) Essentiality features, such as context-dependent tolerance score (CDTS)^23^ and probability of loss-of-function intolerance (pLI)^33^, among others. The latter was used by mapping each non-coding genetic variant to the closest gene and assigning the gene essentiality score of that gene to the corresponding variant. (ii) Chromatin structure features, such as chromosome conformation^23,25^ data used either as a binary indicator to denote whether or not a given non-coding genomic position physically interacts with gene promoters, or as a continuous feature, by attributing the respective gene essentiality of the associated promoter to the distal interacting region. The loop and anchor features were also used as discrete values representing the number of cell lines where they were identified. (iii) Gene expression related features, such as readout of high-throughput enhancer functional screens^26^, and (iv) existing non-coding deleteriousness metrics: CADD^13^, ncEigen^14^, FATHMM^17^, FunSeq2^16^, LINSIGHT^21^, ORION^20^, ReMM^18^ and ncRVIS^29^.

We scored the non-coding regions genome-wide. The result of this process was a score (*ncER*, non-coding Essential Regulation) for each nucleotide, ranging from 0 (non-essential) to 1 (essential). We evaluated the model performance on a test set comprising 20% of the data through 5-fold cross validation and assessed the generalization of the model on 3 independent non-overlapping sets, consisting of curated Mendelian and two hold-out HGMD and ClinVar variants, for validation of the performance of the classifier (**Methods**). The ensemble algorithm, ncER, with a Receiver Operating Characteristic (ROC) AUC of 93% and a Precision-Recall (PR) AUC of 75% on the test set outperformed previously reported deleteriousness metrics by at least 18% ROC-AUC and 31% PR-AUC (**Figures 1A and B**). The model generalized to other independent validation datasets, achieving a ROC-AUC ranging from 88% to 96% and a PR-AUC ranging from 63% to 86% (**Suppl. Figure S2**). The univariate importance of each input feature in the model is displayed in **Figure 1C**. Most of the features (34 out of 38) in the model contributed to the score. The top contributing features were CDTS^23^ that measures human-specific genomic constrain, 3D organization features such as distal enhancers^46^ of essential genes and GTEx^30^ expression variance as indicator of tolerance to gene dysregulation. Because of sparsity of some of the data (for example, pcHi-C and Vista enhancers), some of the features did not contribute to the model because few variants mapped to informative positions. A model trained with only the new essentiality, 3D genome organization and gene expression features outperformed a model trained with only previously published metrics (**Figures 1D and E**). In summary, the increase in ROC AUC and PR-AUC observed after inclusion of new features in the models emphasizes their orthogonality to the previously reported scoring metrics.

**Figure 1.**
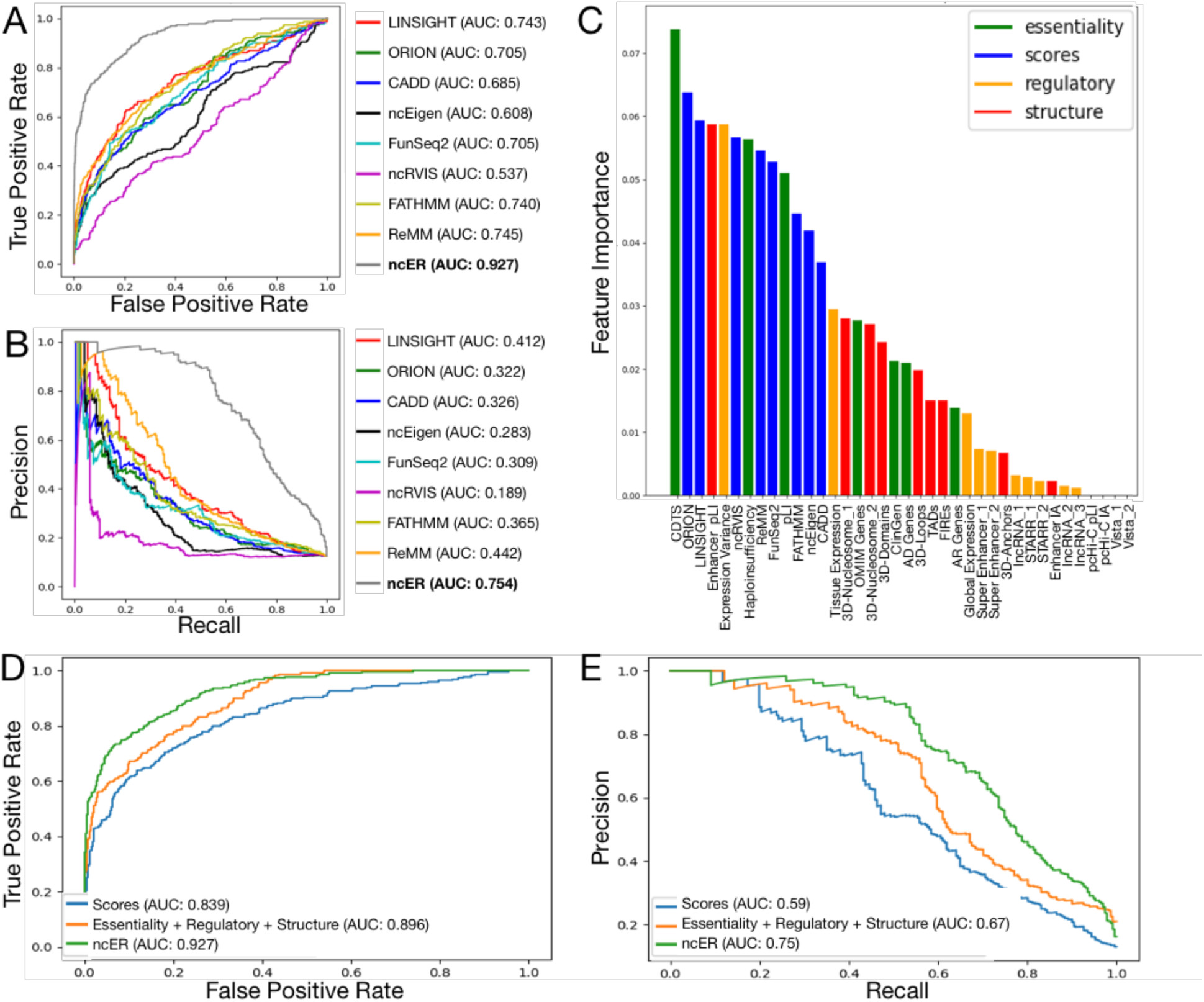
Ensemble learning for the prediction of essential regulatory domains. Performance ROC-AUC (**panel A**) and PR-AUC (**panel B**) on the test set (N=231 non-coding pathogenic and N=1,607 control variants) of ncER model (in grey) compared to previously published scores. The color codes for each metric is shown in the legend. The various input features to ncER (feature importance, the features that have the most effect in the model) are shown in **panel C**. Green, new essentiality features; Blue, published scores; Orange, regulatory/functional screen features; Red, 3D chromatin structure features. Performance ROC-AUC (**panel D**) and PR-AUC (**panel E**) of model trained only with published deleteriousness metrics (blue), only with new features (orange) and with both new features and published metrics (green, ncER).

For computing efficiency purposes and to increase the signal-to-noise ratio, we smoothed the scores over 10bp window and used the 10bp bin resolution for all subsequent analyses. We examined the nature of the most essential regions of the genome based on different ncER percentile thresholds (99.9^th^, 99.5^th^, 99^th^ and 95^th^ percentiles, representing 2.8, 14.1, 28.3 and 141 Mb of cumulative sequence). The distribution of essential genomic elements at each threshold is displayed in **Suppl. Figure S3A**. All types of genomic elements were represented in the most essential bins of the genome, although *cis*-regulatory and enhancer sequences were enriched in the highest percentiles. Essential regions were of small size, with the most common size range being the 10bp bin (**Suppl. Figure S3B**). To have an overview of the putative function of those essential regulatory regions, we did pathway analyses for the set of genes (N=2,441) with at least one promoter bin in the top 99.9% ncER values. The most significant enriched biological process GO terms included development and regulation of gene expression, such as cerebral cortex development, positive/negative regulation of transcription and gene expression, among others (**Suppl. Figure S4**).

In summary, a model that trains on novel genomic features (essentiality, 3D organization, expression) adds precision to previous models that trained on partially orthogonal features (biochemistry, conservation). The model performs well in testing and generalization using new sets of human non-coding disease variants.

## Functional correlates of essential regulatory elements

To confirm that ncER signals effectively captured essential regulatory functions, we analyzed two sets of functional data. The first analysis used high-throughput CRISPRi data from 1.29 Mb of sequence in the vicinity of two essential transcription factor genes, *GATA1* and *MYC*^31^. The library deployed more than 80,000 single guide RNA (sgRNAs) pairs tiled across the genomic loci. The readout of the study was cellular proliferation of K562 erythroleukemia cells. *GATA1* or *MYC* regions are characterized by a high median ncER score, in the 87th percentile (**Figure 2A and 2B** and **Suppl. Figure S5**) – which is consistent with the biological importance of the loci. However, we observed a shift to a median 99th ncER percentile for the regions targeted by the most functional pairs of sgRNA probes (p=2.1e-15, compared to non-functional probes). There was a dosage effect: the more essential the region, the stronger the functional readout of CRISPRi (**Figure 2B**). The parameters of predictive performance and accuracy of ncER are shown in **Suppl. Table S2**. In summary, the sequences that control cellular proliferation, cell viability and gene expression of *GATA1* and *MYC* reside in the most essential regions of the locus.

**Figure 2.**
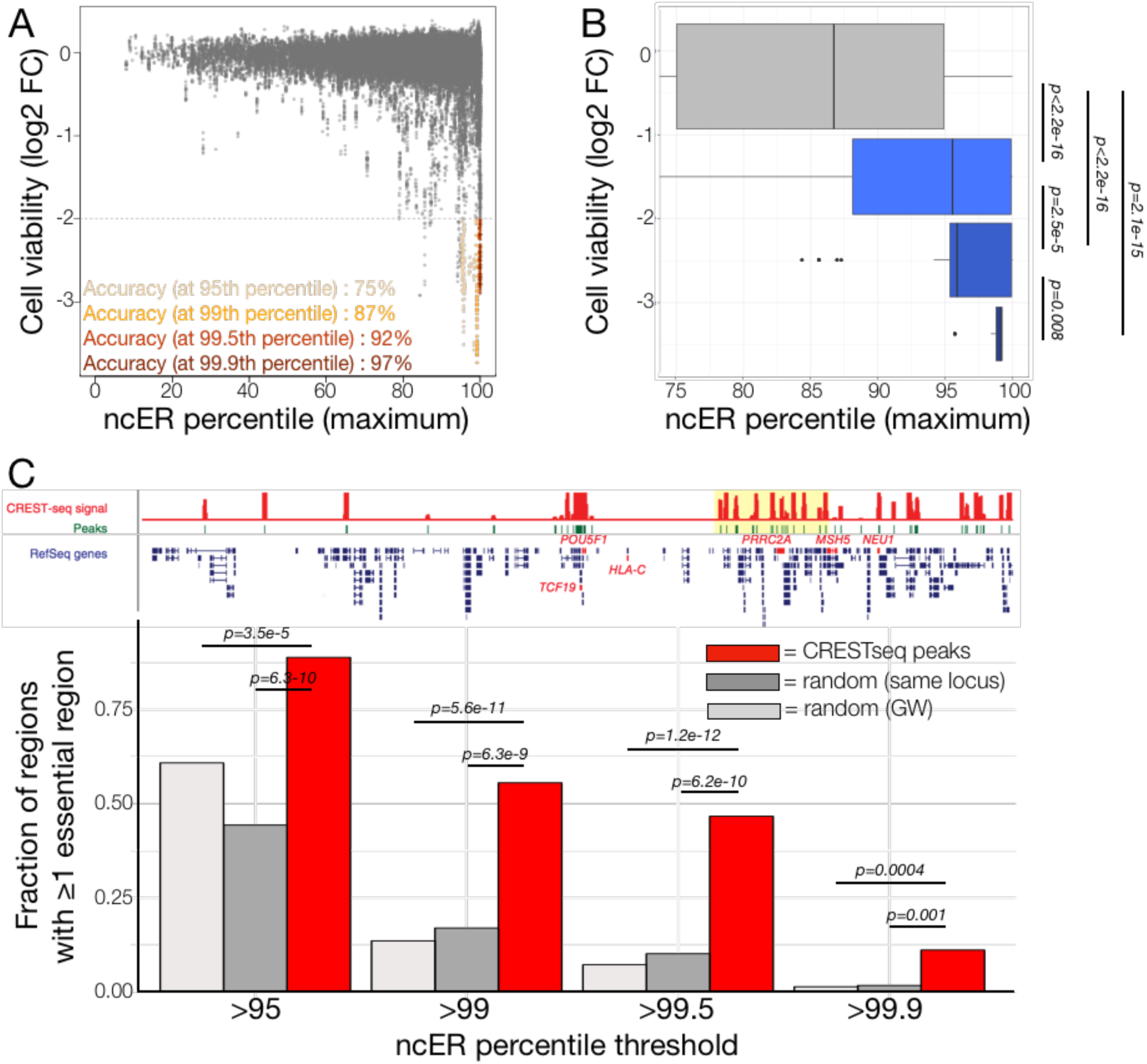
Comparison of experimental functional assays with *in silico* predictions of essentiality. **Panel A.** CRISPRi effect on cell viability (77,368 sgRNA probes pairs) from Fulco et al.^31^ and the corresponding maximum ncER score accross the *GATA1* and *MYC* loci. Accuracy at four ncER thresholds is shown in yellow, orange, red and dark-red respectively for the 95^th^, 99^th^, 99.5^th^ and 99.9^th^ percentiles. **Panel B.** Distribution of maximum ncER at different bins of cell viability (0 to less than -3 log2 fold change). P values were computed with independent 2-group Man-Whitney Unpaired Test. **Panel C.** Upper display represents the experimental locus and the identified CRESTseq peaks (n=45, green), from Diao et al.^32^. Lower display presents the fraction of regions with at least one essential bin. Essential bins are defined by four different ncER percentile thresholds. CRESTseq peaks are shown in red (N=45), random matched sized *in silico* regions extracted from the same locus (“random same locus, N=4,500) in dark grey and random matched sized *in silico* regions extracted genome-wide (“random GW”, N=4,500), in light grey. P values were computed with Fisher Exact Test. Panel C pictogram is adapted from^32^. FC, fold change. CRESTseq, *cis*-regulatory elements by tiling-deletion and sequencing. GW, genome-wide.

The second dataset was generated using high-throughput scanning for *cis*-regulatory elements by tiling-deletion and sequencing (CREST-seq)^32^. The area investigated encompassed 2-Mb of the *POU5F1* locus in human embryonic stem cells. *POU5F1* encodes a transcription factor that plays a key role in embryonic development and stem cell pluripotency. Knockout of *POU5F1* is associated with embryonic mortality in the mouse and scores as an essential gene in humans (pLI score of 0.89)^8,33^. Thus, *POU5F1* is expected to use regulatory elements with features of essentiality^23^. CREST-seq identified 45 *cis*-regulatory elements, including 17 previously annotated as promoters of unrelated genes that, like typical enhancers, form extensive spatial contacts with the *POU5F1* promoter. The 45 enhancers of *POUSF1* were significantly more likely to be essential compared to random genomic loci of matched size (**Figure 2C**). For example, 56% of *POU5F1* enhancers reside in regions with ncER >99^th^ percentile, compared to 14-17% for random genomic regions matched to enhancer size, p≤6.3e-09. Similarly, the enhancers contained the most essential regions within the locus in permutation analysis (**Suppl. Figure S6**). The parameters of predictive performance and accuracy of ncER are shown in **Suppl. Table S3** and **Suppl. Figure S7**.

In summary, the excellent performance of ncER scores for the identification of deleterious variants and for the prediction of functional read-outs in the non-coding genome may help map critical regulatory and structural elements of the non-coding genome in human disease.

## Mapping essential regulatory elements in genetic diseases

We hypothesized that severe genetic diseases that do not have causal variants in the coding region could result from damage to essential functional elements. To investigate this concept, we chose two different models that represent challenges for the accurate mapping of the regulatory sites in relation to function and disease: (i) the identification of essential functional areas within non-coding structural variants/deletions associated with autism and ASD, and (ii) the impact of reorganization of TADs in the setting of a human developmental disorder that has been modeled in the mouse.

For the first disease model and as follow-up to our previous collaboration with Sebat et al.^34^, we assessed a dataset of 136 transmitted *cis*-regulatory structural variants (SV, deletions) in the setting of autism and ASD. The deletions spanned a wide range of sizes: <1kb, N=32 probands and N=10 healthy sibling controls; 1-25kb, N=65 cases and N=4 controls; 25-100kb, N=17 cases and N=2 controls. The last group, >100kb deletions, found in N=6 probands did not have matched controls (**Suppl. Figure S8**). We observed that probands carry structural variants with only slightly higher ncER scores (median ncER percentile: 86 in probands versus 70 in healthy siblings for the <1kb deletions, p=0.21, **Suppl. Figure S9**). However, autism and ASD probands were more likely to carry structural variants with localized essential functional domains compared to healthy siblings (**Figure 3A**). For example, 41% of the <1kb deletions in probands contain regions with ncER >99^th^ percentile, compared to 4% for random genomic regions matched to the size of the deletions, p=1.7e-10 and also compared to the few deletions present in non-probands (**Figure 3A**). Similar findings were observed for larger deletions (**Suppl. Figure S10**). As example, 20 unique <1kb deletions with at least one essential domain are displayed in **Figure 3B**. These results indicate that the identification of essential regulatory regions could map the most plausible causal region within a structural variant.

**Figure 3.**
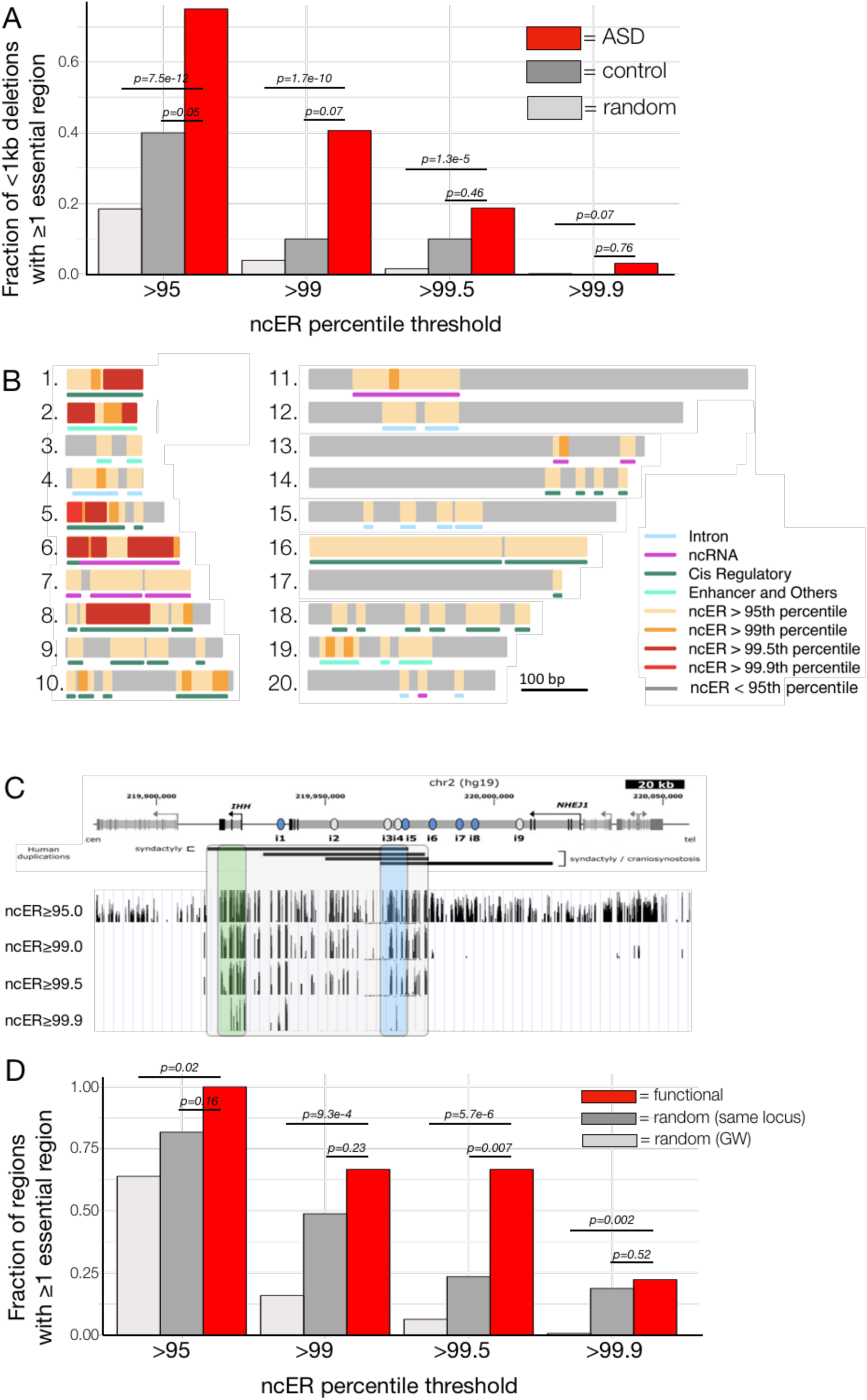
Mapping of essential regulatory domains in disease models. **Panel A.** Fraction of <1kb deletions with at least one essential bin. Essential bins are defined at four different ncER percentile thresholds. Autism and ASD deletions are shown in red (ASD, N=32), control deletions in dark grey (control, N=10) and random size-matched *in silico* deletions extracted genome-wide in light grey (random, N=4,200). See **Suppl. Figure 8** for distribution of other sizes of deletions. **Panel B**. Schematic illustration of 20 unique deletions (grey bars) identified in Autism and ASD probands that harbor essential regions (highlighted in yellow, orange, red and dark red, based on the ncER thresholds). The corresponding genomic elements present at the essential regions are displayed under the deletions. Introns are shown in blue, ncRNA in magenta, *cis* regulatory in dark green and enhancers and others regulatory elements in light green (See **Methods** for categorization of genomic element classes). **Panel C**. The upper panel illustrates the human *IHH* genomic locus associated with developmental defects including craniosynostosis and synpolydactyly^35,36^. It harbors the 9 enhancers identified in mice (from Will et al.^37^). Lower panel, UCSC genome browser view of the essential region in the locus. Essential bins are shown at four ncER percentile thresholds. The grey box inset highlights the region of essentiality across human pathogenic duplications, the blue box the essentiality at the maximal overlap of genomic lesions in humans, and the green box that includes the *IHH* region present in duplications causing syndactyly Leuken type engineered in Will et al.^37^. Panel C pictogram is adapted from^37^. **Panel D.** Fraction of regions with at least one essential bin. Essential bins are defined by four different ncER percentile thresholds. Mouse to human mapped enhancers are shown in red (“functional”, N=9), random size-matched *in silico* deletions extracted from the same locus (“random same locus”, N=900) in dark grey and random size-matched *in silico* deletions extracted genome-wide (“random GW”, N=900) in light grey. GW, genome-wide.

Next, we chose a human disease that involves the rearrangement of the regulatory landscape of *IHH* (encoding Indian hedgehog), which cause developmental defects including craniosynostosis and synpolydactyly^35,36^. Will et al.^37^ identified nine enhancers with individual tissue specificities in the digit anlagen, growth plates, skull sutures and fingertips. The *IHH* region in humans shares a common structure with the mouse locus that was used for the model by Will et al.^37^ In their study, consecutive deletions resulted in growth defects of the skull and long bones that confirmed that the enhancers function in an additive manner. Deletions and duplications caused dose-dependent upregulation and misexpression of *Ihh*, leading to abnormal phalanges, fusion of sutures and syndactyly. We identified the critical enhancers to reside in an extensive region of essentiality as scored by ncER, in particular for the regions shared across human duplications associated with disease (**Figure 3C**, Blue box). Within the locus, the critical enhancers were also endowed with features of essentiality (**Figure 3D**). For example, 67% of the enhancers contain regions with ncER >99^th^ percentile, compared to 16% for random genomic regions matched to size, p=9.3e-04. Similarly, the nine enhancers carried the most essential regions within the locus in permutation analysis (**Supp. Figure S11**). Thus, highly essential enhancers that associate with human disease are correctly prioritized by computational approaches.

The present work contributes to the debate about reconciling redundancy and conservation of regulatory elements in the genome. Work on developmentally expressed genes supports the widespread existence of functionally redundant enhancers in mammalian genomes^38^. Redundancy reduces the likelihood of severe consequences resulting from genetic or environmental challenge. However, Osterwalder et al.^38^ also suggest that contributions of enhancers to overall gene expression levels are relevant for organismal fitness under specific pressures, thus subjecting enhancers to purifying selection over evolutionary time. We have in the past indicated that essential genes will use proximal and distant regulatory elements that are conserved and constrained – thus representing essentiality in the non-coding genome^8,23^. We now show hallmarks of proximal or distal regions that regulate the expression of medically important genes. The current model supports the prioritization of variants and regions across the non-protein coding human genome for diagnostics and for functional analysis.

## Materials and Methods

### Training Features

To train our model, we leveraged a total of 38 features from 4 major categories: (i) gene essentiality, (ii) 3-dimentional chromatin structure, (iii) gene expression and other regulatory/functional data and (iv) existing variant pathogenicity/deleteriousness scores. A complete list of features along with their descriptions and accession links can be found in **Suppl. Table S1**.

We have previously identified a coordination of constrains between genes and their respective *cis* and distal regulatory elements^23^. We implement this concept in the present study by including the following essentiality features: (i) CDTS (our recently developed approach to score the non-coding genome essentiality, based on human genetic diversity^23^, (ii) probability of loss-of-function intolerance (pLI)^33^, (iii) haploinsufficiency score^39^, (iv) gene dosage sensitivity score from ClinGen^40^, (v) autosomal dominant or recessive categorization^41,42^ and (vi) OMIM association ^43^. For the metrics that solely provide scores for the genic portion of the genome, the respective essentiality features were calculated by mapping each non-coding genomic position to the nearest gene and assigning the corresponding essentiality metric score to the genomic position.

Chromatin 3D structure features included (i) nucleosome positioning extracted from MNase data (https://www.encodeproject.org/), (ii) multiple cell type anchor, loop and domain regions extracted from Hi-C data^44^, (iii) frequently interacting regions (FIRE) and topologically associated domains (TADs) extracted from Schmitt et al.^45^ and (iv) distal enhancer-TSS associations extracted from CAGE pairwise expression correlation (FANTOM)^46^. The 3D organization features were either used as binary indicators to denote whether or not a given non-coding genomic position physically interacted with gene promoters, or as discrete values representing the number of cell lines were the structures were identified. Finally, to combine both essentiality and chromatin structure features, we created distal essentiality features, by attributing the respective coding gene essentiality score (pLI) to distal regulatory elements identified through pcHi-C or CAGE pairwise expression correlations.

The model used gene expression, long non-coding RNA (lncRNA) annotations and functional regulatory data that have not been used by other existing metrics. Those included (i) median gene expression and variance across tissues (GTEx)^30^, (ii) functionally tested genomic regions with enhancer activity identified through (ChIP-)STARR-seq experiments^26,47^ or validated with transgenic mice^48^ and (iii) lncRNAs identified through CAGE and transcriptome analysis^49^.

Lastly, variant pathogenicity/deleteriousness scores used in the model included CADD^13^, ncEigen^14^, FATHMM^17^, FunSeq2^16^, LINSIGHT^21^, ncRVIS^29^, Orion^20^ and ReMM^18^. We downloaded pre-computed genome-wide scores for each of these metrics (hg19 reference build). In the minority of cases where a per alternative variant score was provided, we used the most “deleterious” value at each position. For the metrics that solely provide scores for the genic portion of the genome, the respective features were calculated by mapping each non-coding genomic position to the nearest gene and assigning the corresponding metric score to the genomic position.

### Training Set

We used a total of 9,188 single nucleotide variants (SNVs) to train the model. The pathogenic dataset comprised 1,095 non-coding SNVs, located at least 10bp from the nearest spice site, obtained from HGMD (2016_R1)^28^ and ClinVar (July 2016)^27^. The selection criteria for HGMD SNVs were “DM and high” tags, while for ClinVar, SNVs had to be labeled as “Pathogenic” or “Likely Pathogenic”, with star 1 or more and no conflicting assertion. HGMD were further filtered out for variants overlapping SNVs annotated as “benign” or “likely benign” in ClinVar (with star 1 or more and no conflicting assertion). The control genomic training set consisted of 8,093 variants chosen from a larger set of variants that were present in the gnomAD dataset at an allelic frequency > 1% and matched for the distance to the nearest splice sites and genomic elements. The matching was performed as follows: all pathogenic variants and gnomAD variants with allelic frequency > 1% were annotated with their respective distance to the closest splice site and the genomic element they mapped to (see Reference build, Annotation and Genomic Element Categorization section). For each pathogenic variant the subset of control variants falling within the same genomic element was extracted. Within this subset, the 10 control variants with the most similar distance to splice site as compared to the pathogenic variant were kept. Finally, duplicated control variants (if any) were removed from the final set. 80/20 percent of the variants were respectively used as training/test sets.

### Machine Learning Model

We trained an XGBoost model in order to differentiate between pathogenic and control genomic positions in our training set. Hyperparameters were tuned using 5-fold cross validation and a randomized search method. 1,000 sets of randomly selected hyperparameters were evaluated using 5-fold cross validation, and the model that achieved the highest ROC-AUC score was selected. These hyperparameters were then used to train the final model on the entirety of the training set. After hyperparameter tuning, we found that using 233 estimators, a maximum depth of 31, a learning rate of 6.1e-2, and a minimum child weight of 6.17 maximized model performance. We evaluated our model with Receiver Operating Characteristic (ROC) AUC and Precision-Recall (PR) AUC on the test set (representing 20% of the data).

We annotated each position in the genome with our set of features and used the tuned XGBoost model to make a functionality prediction at each genomic position to score the entire genome. Of note, the model is trained to assess the functionality and essentiality of regulatory regions and should therefore be interpreted as such even in protein coding regions.

### Validation Sets

The generalization of the model was assessed on three independent sets of variants. The non-coding pathogenic sets included 425 manually curated variants associated with Mendelian traits^23^, 354 and 245 new HGMD and ClinVar variants^28^ mapping outside/inside non-coding RNA genes (HGMD 2017_R2 and ClinVar January 2018). The control genomic training sets consisted respectively of 1,863, 2,604 and 1,876 variants in the gnomAD dataset at an allelic frequency > 1% and matched to the pathogenic sets for the distance to the nearest splice sites and genomic elements as explained above.

### Reference build, Annotation and Genomic Element Categorization

All input features and the model were mapped to the human reference build hg19. To investigate the element distribution, we built an annotation track that combined annotations from GenCode (v.27 mapped to GRCh37) and ENCODE (annotated features and multicell regulatory elements, Ensembl v91 Regulatory Build) and used a prioritization scheme to assign each genomic position a single annotation category (described in ^23^). For **Figure 3** and **Suppl. Figure S3**, *Intron* refers to intronic regions (from protein coding or non-coding genes), *ncRNA* refers to exonic regions of non-coding RNAs, *Cis Regulatory* encompasses promoters and untranslated regions (UTRs), *Enhancers and Others* encompasses promoter flanking regions, enhancers, open chromatin, CTCF and other transcription factor binding sites, *Intergenic* refers to unannotated regions and *Histone marks* encompasses H3K9me3 and/or H3K27me3 as well as other histone marks combinations.

### ncER score

For computing power purposes and to minimize the signal-to-noise ratio, the ncER score were averaged over 10bp bins and then expressed as percentiles genome-wide. The binned/percentile scores are provided at https://github.com/TelentiLab and can be accessed directly at OMNI (https://omni-variants.herokuapp.com/). Intersection of ncER score with other datasets was performed using bedops utility (v2.4.30)^50^.

### External Datasets

The CRISPRi coordinates and scores used for analyses displayed in **Figures 2A and 2B** and **Suppl. Figure S5** were obtained from Fulco et al.^31^ (http://science.sciencemag.org/highwire/filestream/686019/field_highwire_adjunct_files/2/aag2445_Table_S2.xlsx). As recommended in their paper, the CRISPRi scores were smoothed over 20 subsequent pairs. . Only regions that were assessed by at least 20 different pairs and <1kb long (to prevent size biases) were retained for analysis, resulting in a total of N=77,368 remaining probed pairs.

CREST-seq peaks coordinates used for analyses displayed in **Figure 2C** and **Suppl. Figure S6** were obtained from Diao et al.^32^ (https://media.nature.com/original/nature-assets/nmeth/journal/v14/n6/extref/nmeth.4264-S7.xlsx). Statistical enrichment of the 11,570 tested sgRNA pairs used for analyses in **Suppl. Figure S7** were also obtained from 32 (https://media.nature.com/original/nature-assets/nmeth/journal/v14/n6/extref/nmeth.4264-S5.xlsx). The locus used for random extraction of same size regions was chr6:30132133-32138339. The matched size random extraction (both in the same locus and genome-wide) was performed 100 times.

*Cis* regulatory transmitted deletions used for analyses displayed in **Figures 3A and B** and **Suppl. Figures S8-10** were obtained from Brandler et al.^34^ (http://science.sciencemag.org/highwire/filestream/708877/field_highwire_adjunct_files/9/aan2261_TableS7.xlsx, Replication CR Trans sheet). The matched size genome-wide random extraction was performed 100 times.

The nine mouse enhancer data used for analyses displayed in **Figure 3C** and **Suppl. Figure S11** were obtained from Will et al.^37^ (https://media.nature.com/original/nature-assets/ng/journal/v49/n10/extref/ng.3939-S1.pdf, Supplementary Table S4). The mouse coordinates were mapped to human using CrossMap (v.0.2.5. http://crossmap.sourceforge.net/) using the mm9ToHg19.over.chain.gz chain. When the mouse enhancers were mapped discontinuously to the human genome, the left most and right most coordinates in the human genome were used as start and end. The locus used for random extraction of same size regions was chr2: 219940039-220025587. The matched size random extraction (both in the same locus and genome-wide) was performed 100 times.

### Statistics

Statistical analyses and plotting were performed with R v3.4.3 (https://www.R-project.org/), notably using the package ggplot2 (http://ggplot2.org/). Data mining was performed using Python (v.2.7.11). The performance predictors in **Figures 1** and **2, Suppl. Figures 2,5** and **7**, **Suppl. Tables S2** and **S3** were assessed as follows: sensitivity or true positive rate or recall is (TP/(TP+FN)) * 100, specificity is (TN/(TN+FP)) * 100, false positive rate is (FP/(FP + TN)) * 100, accuracy is ((TP+TN)/(TP+TN+FP+FN))*100, positive predictive value or precision is (TP/(TP+FP))*100 and negative predictive value is (TN/(TN+FN))*100. Where TP is a true positive, TN is a true negative, FP is a false positive and FN is a false negative.

## Acknowledgments

We thank J. Fellay for comments on the manuscript. Work of A.Te. is supported by the Qualcomm Foundation and the NIH Center for Translational Science Award (CTSA, grant number UL1TR002550).

## Author contributions

Conception and design of the study: J.d.I, A.Te. Performed the analyses: A.W., J.d.I. Built the browser and code repositories: L.Y. Contributed methods and analytical strategies: D.H., A.To., B.R. Wrote the manuscript: J.d.I., A.Te.

## Supplementary materials

**Suppl. Figure S1.**
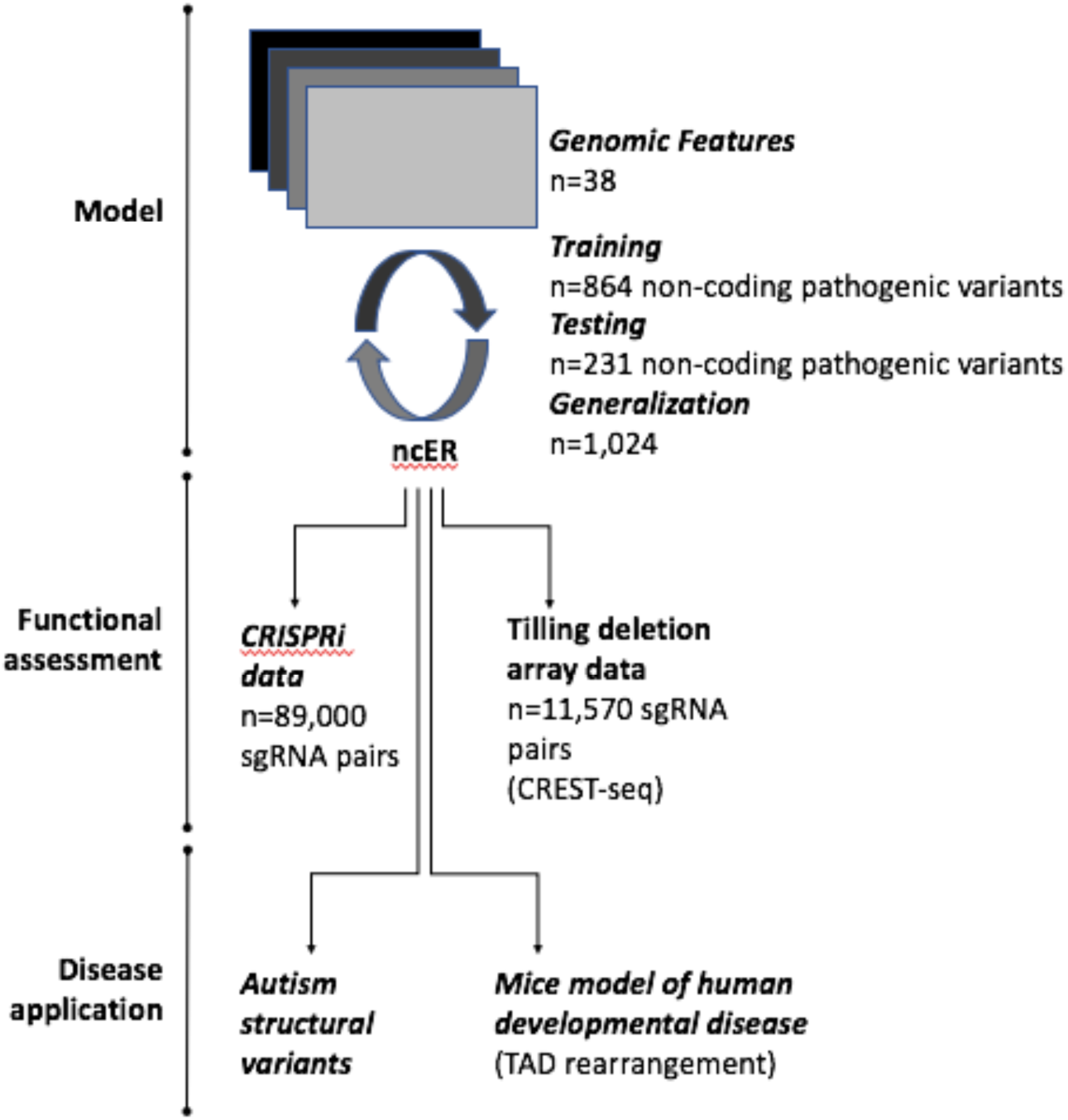
Study design. CREST-seq= *cis-*regulatory element scan by tiling-deletion and sequencing. CRISPRi= clustered regularly interspaced short palindromic repeats (CRISPR) interference.

**Suppl. Figure S2.**
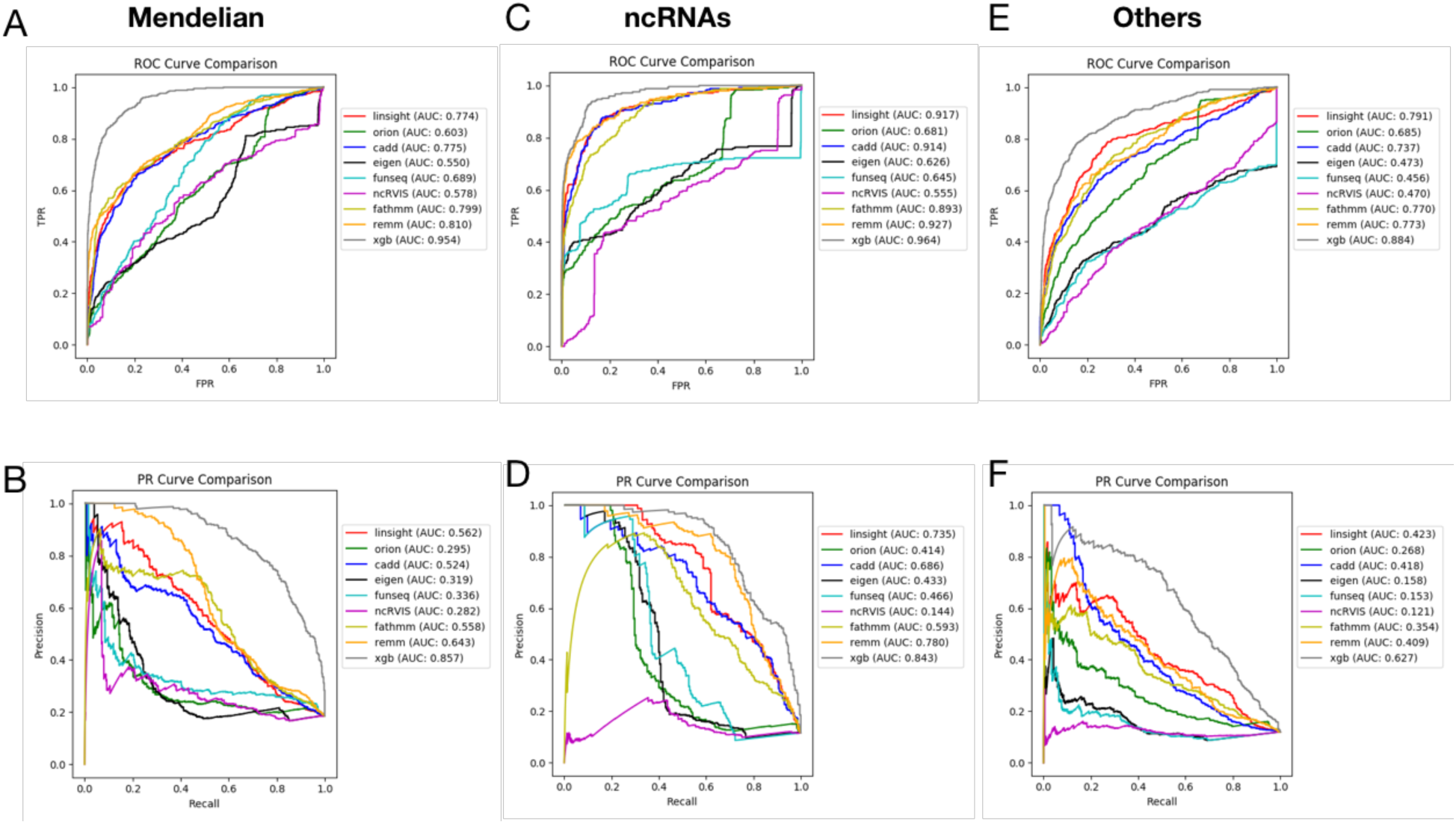
ncER generalization to independent variant sets. Performance ROC-AUC (**panel A**) and PR-AUC (**panel B**) for the independent set of manually curated non-coding Mendelian variants (N=425 pathogenic and N=1,863 control)^23^. Performance ROC-AUC (**panel C**) and PR-AUC (**panel D**) for the independent set of left out non-coding HGMD and ClinVar variants mapping to ncRNAs (N=245 pathogenic and N=1,876 control). Performance ROC-AUC (**panel E**) and PR-AUC (**panel F**) for the independent set of left out non-coding HGMD and ClinVar variants mapping outside of ncRNAs (Others; N=354 pathogenic and N=2,604 control). ncRNA, non-coding RNA. TPR, true positive rate. FPR, false positive rate. Xgb, XGBoost model or ncER.

**Suppl. Figure S3.**
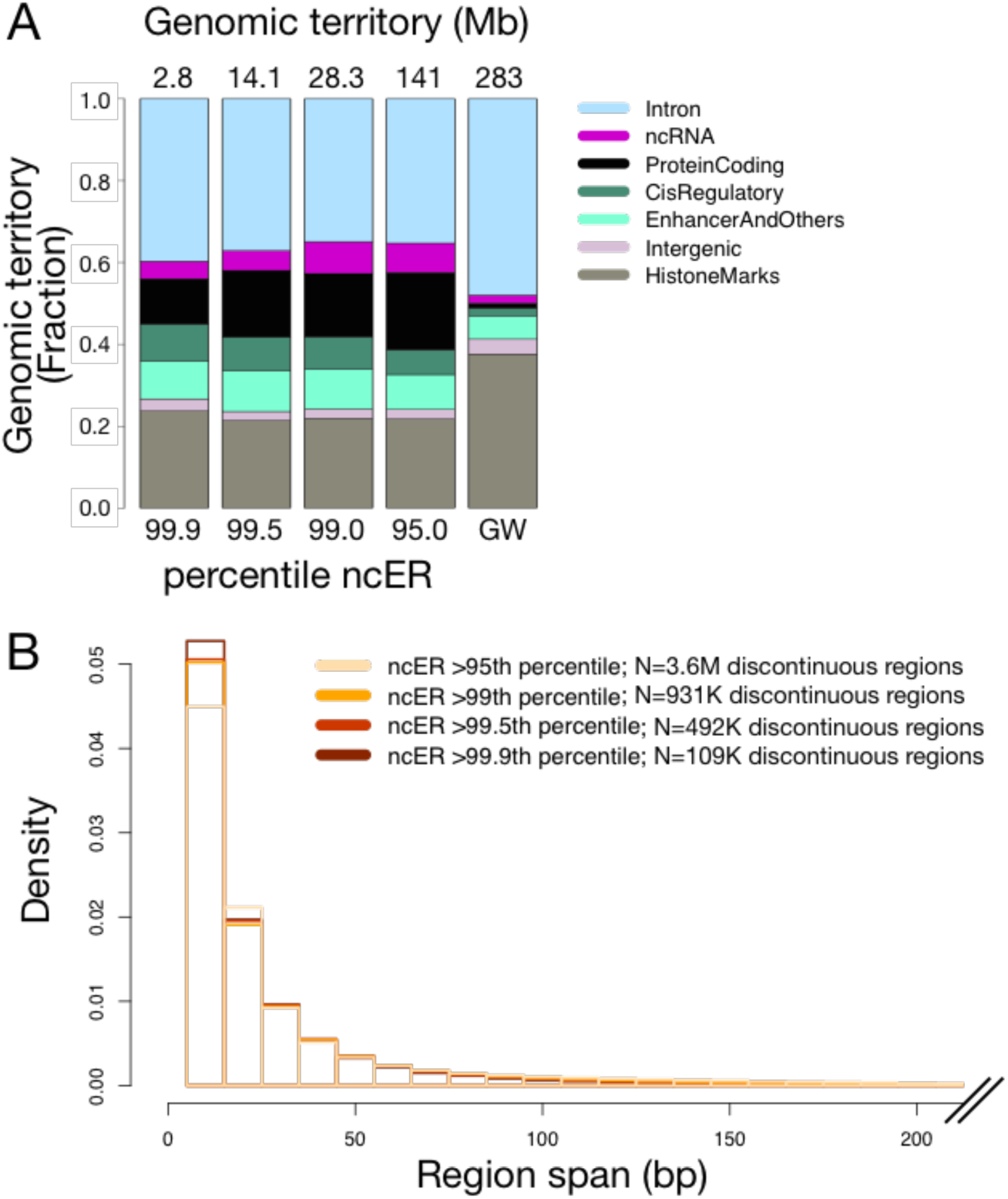
Essential regions distribution in the genome. **Panel A**. Bar plot displaying the cumulative territory covered by each element family at different percentile threshold (indicated at the bottom of the bars). The total genomic territory is displayed at the top of the bars. The percentiles are based on the rank of ncER values. The elements appear in the bar plot in the same order as in the legend. Genomic elements grouping is detailed in **Methods**. **Panel B**. Size distribution of essential regions defined at different ncER percentile thresholds (indicated by the color code in the top-right of the figure). GW, genome-wide.

**Suppl. Figure S4.**
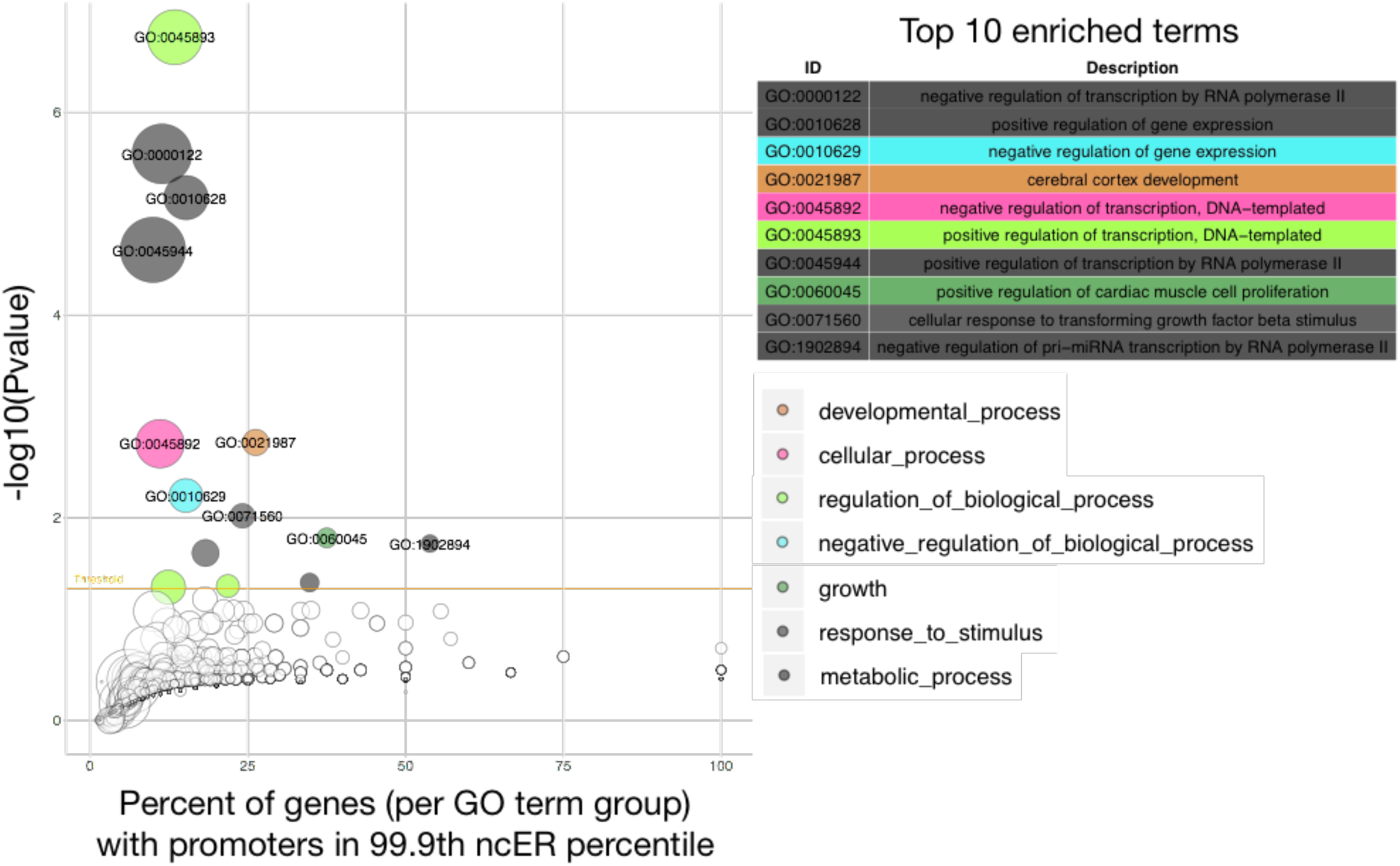
The GO terms enrichment for biological processes in essential regions. The bubble plot represents the significance of the enrichment for a given term (y axis, minus log10(pvalue)) versus the percentage of genes associated with a given biological process that were present in the set of genes with at least one essential promoter bin (x axis). Each circle in the plot represents one biological process. The size of the circles is proportional to the number of genes in the specific GO term class. Only the significant terms are colored. The circles and the rows in the associated table are colored according to the highest ranking hierarchical term (to facilitate pathway and redundant information detection). When multiple terms could be the highest hierarchical ancestor, the coloring was randomly selected among the multiple possibilities. The associated table provides the name of the top ranked terms. Promoter regions were defined as the 600bp upstream the transcription start site. A modified version of the GOBubble function from the GOplot R package (http://wencke.github.io/) was used to generate this figure.

**Suppl. Figure S5.**
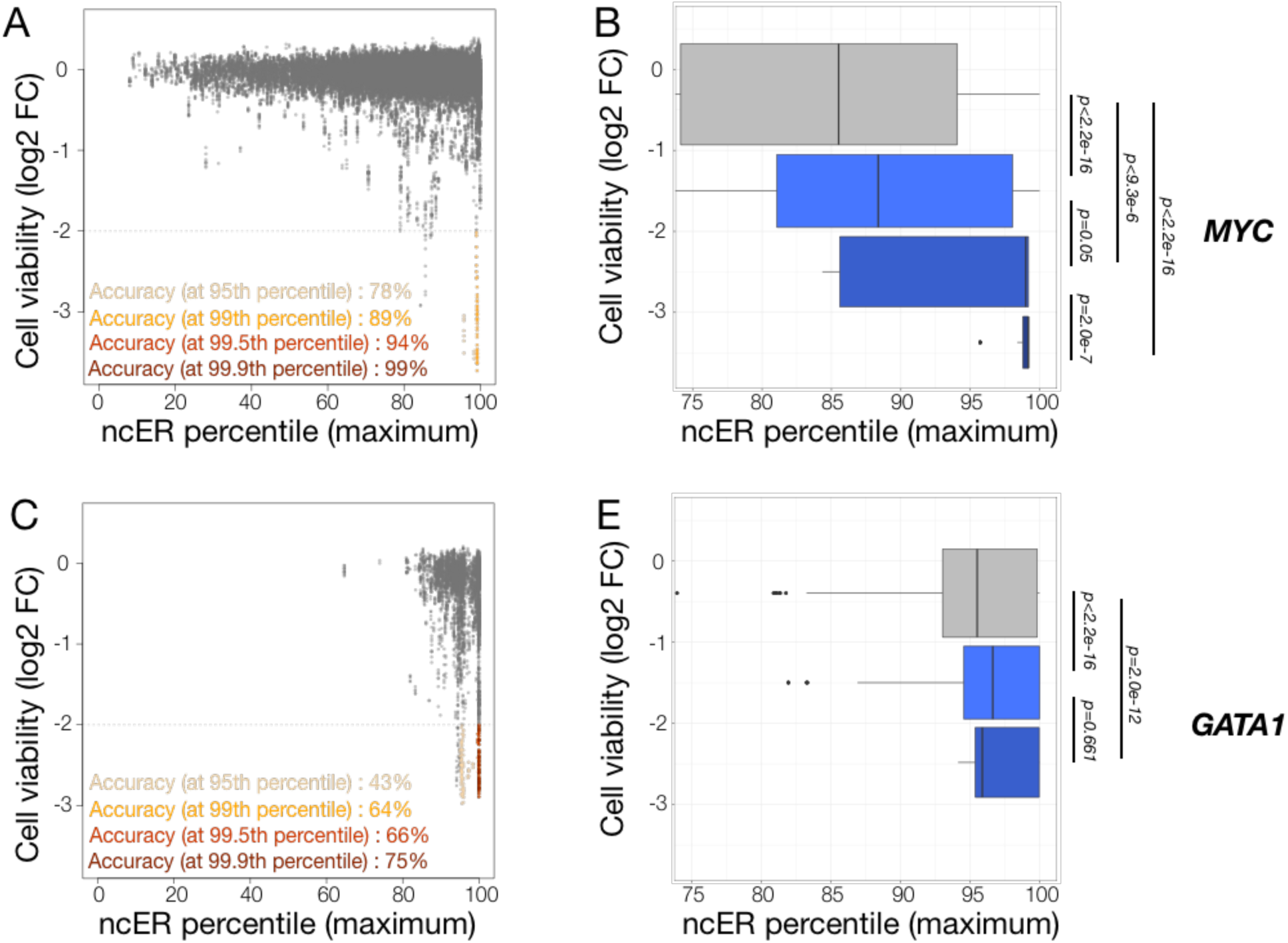
Comparison of experimental CRISPRi functional assays with *in silico* predictions of essentiality. **Panel A.** CRISPRi effect on cell viability (71,404 sgRNA probe pairs targeting the *MYC* locus) and the corresponding maximum ncER score within the tested region. Accuracy at four ncER thresholds is shown in yellow, orange, red and dark-red respectively for the 95^th^, 99^th^, 99.5^th^ and 99.9^th^ ncER percentiles. **Panel B.** Distribution of maximum ncER at different bins of cell viability (0 to lower than -3 log2 fold change). P values were computed with independent 2-group Man-Whitney Unpaired Test. **Panel A.** CRISPRi effect on cell viability (5,856 sgRNA probe pairs targeting the *GATA1* locus) and the corresponding maximum ncER score within the tested region. Accuracy at four ncER thresholds is shown in yellow, orange, red and dark-red respectively for the 95^th^, 99^th^, 99.5^th^ and 99.9^th^ ncER percentiles. **Panel B.** Respective distribution of maximum ncER at different bins of cell viability (0 to lower than -3 log2 fold change). P values were computed with independent 2-group Man-Whitney Unpaired Test.

**Suppl. Figure S6.**
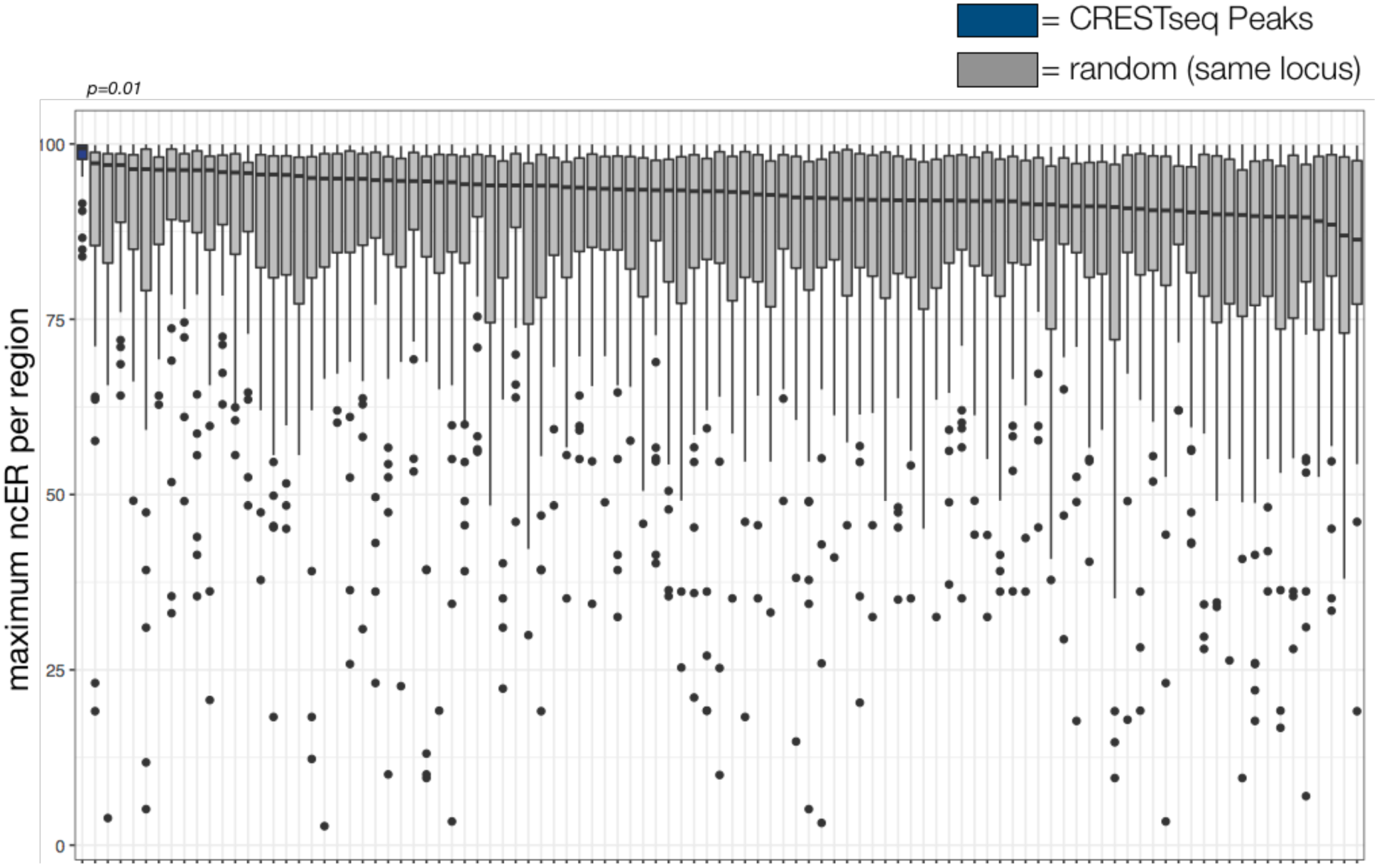
CREST-seq peaks enrichment in essential regions. CREST-seq peaks (N=45, dark blue) display the highest ncER percentile distribution, compared to 100 permutations (grey), each containing 45 regions matched by size to the CREST-seq peaks and from the same genomic locus.

**Suppl. Figure S7.**
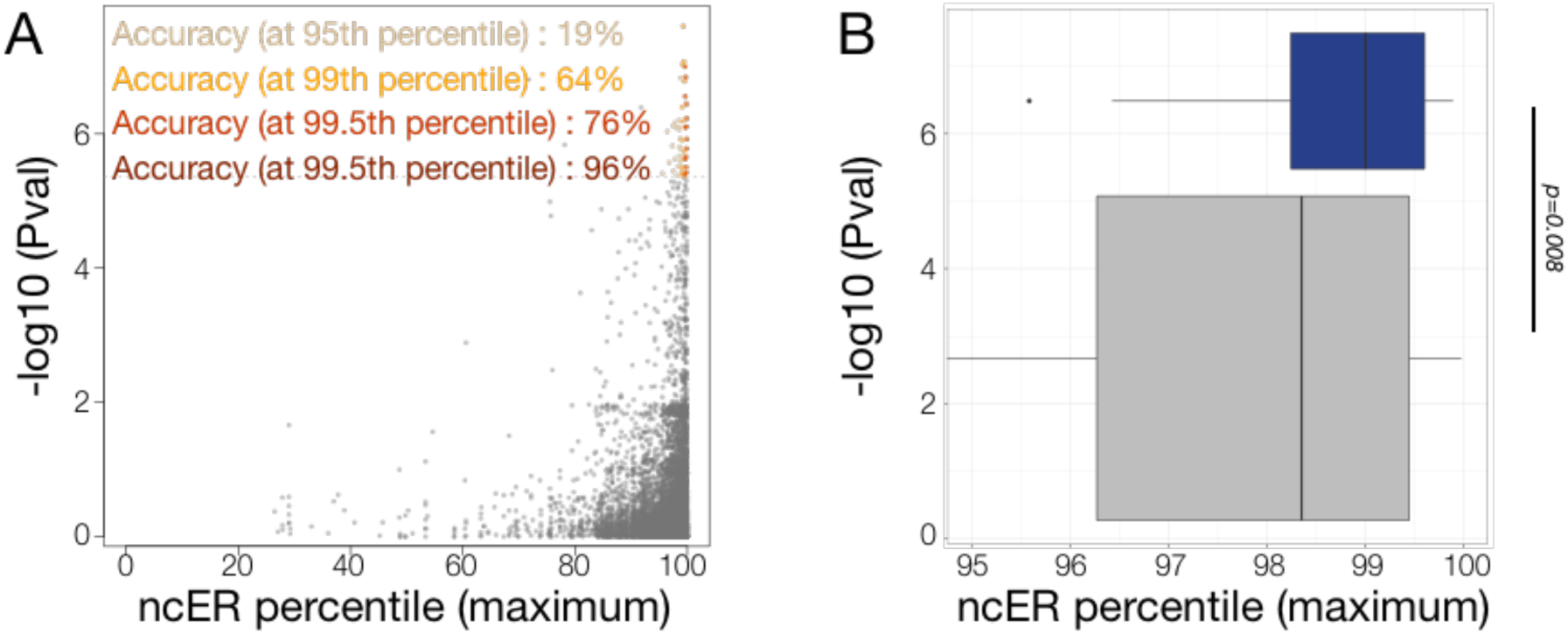
Comparison of experimental CREST-seq functional assays with *in silico* predictions of essentiality. **Panel A.** P values determined by Diao et al. ^32^ comparing *POU5F1* expression in targeted cells versus controls (11,570 sgRNA probe pairs) and the corresponding maximum ncER score within the tested region. Accuracy at four ncER thresholds is shown in yellow, orange, red and darkred respectively for the 95^th^, 99^th^, 99.5^th^ and 99.9^th^ ncER percentiles. **Panel B.** Distribution of maximum ncER at different bins of −log10(p value) (up to 5.36 (non-significant), above 5.36 which corresponds to −log10(0.05/11,570)). P values were computed with independent 2-group Man-Whitney Unpaired Test.

**Suppl. Figure S8.**
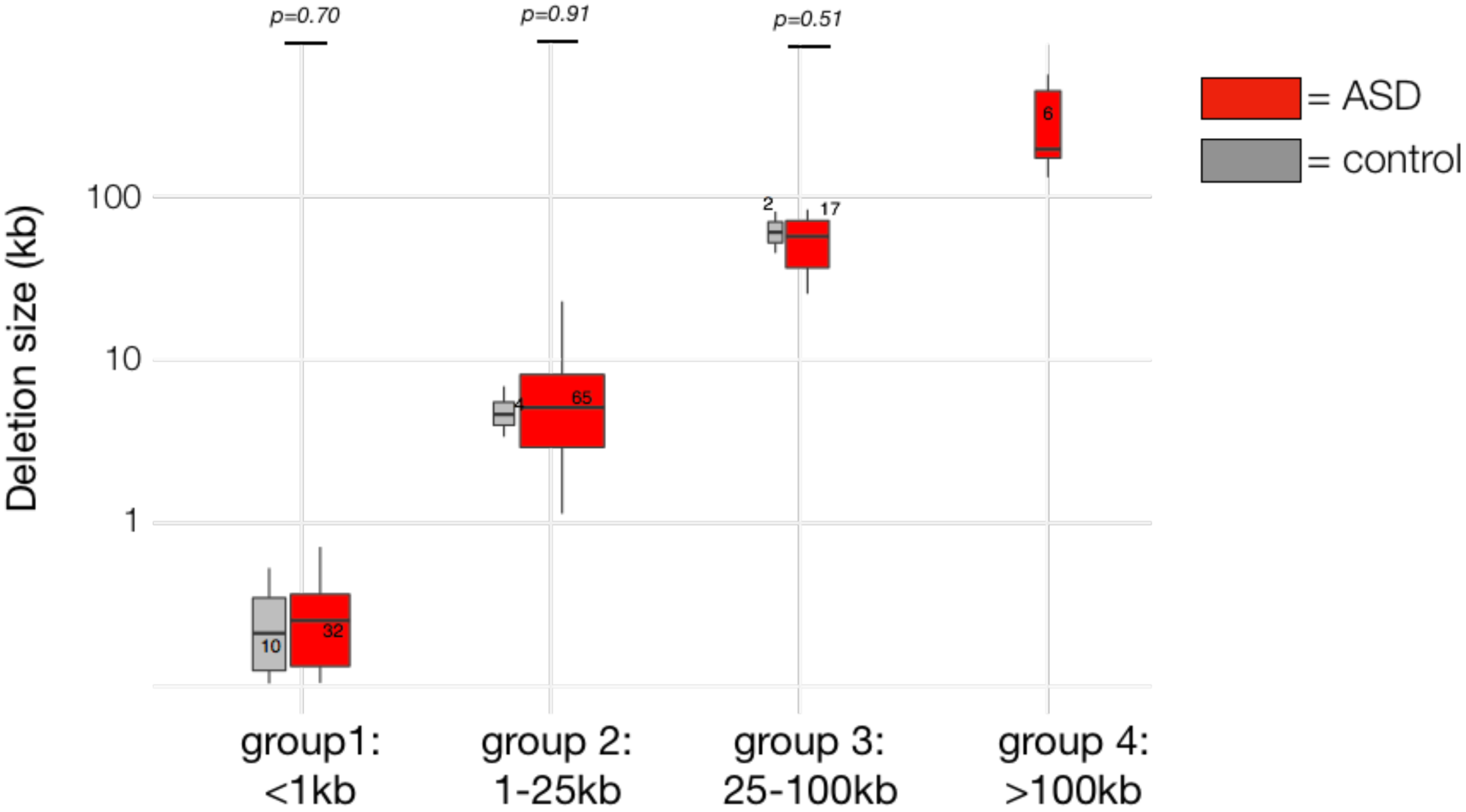
Size distribution of Cis-regulatory transmitted deletions. The transmitted cis-regulatory deletions were split into 4 groups of matched size (with no significant difference in the distribution of control and ASD deletion within the same group size). Deletions present in cases are shown in red, deletions present in controls are shown in grey. The y axis is on logarithmic scale. P values were computed with independent 2-group Man-Whitney Unpaired Test.

**Suppl. Figure S9.**
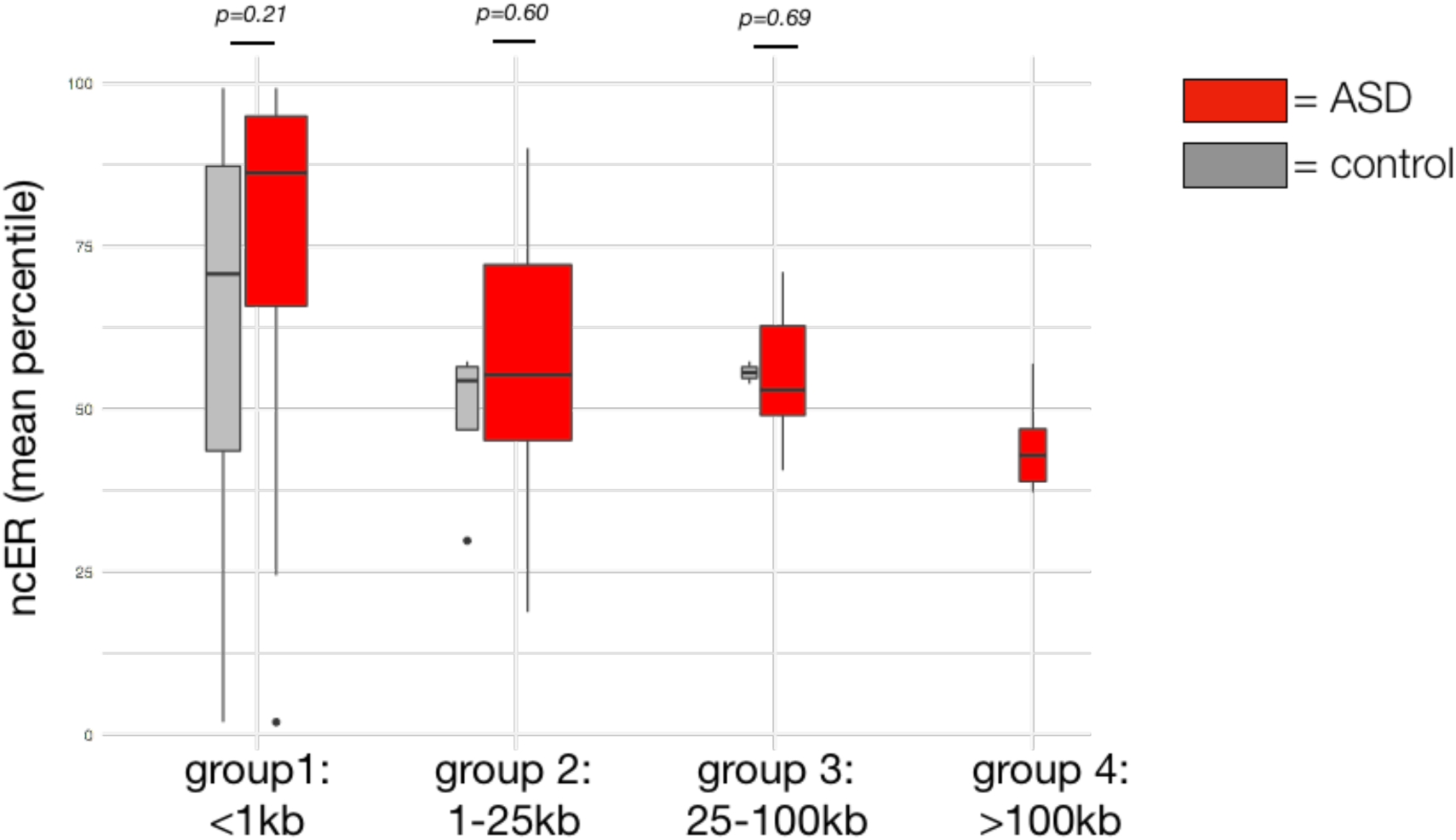
ncER percentile distribution across transmitted deletion. The <1kb cis-regulatory transmitted deletions tend to have an average ncER slightly higher in ASD than in controls, however the effect, if any, disappears for higher size deletions. The longer the deletion the more likely to approach ncER 50^th^ percentile, the genome average. Deletions present in probands are shown in red, deletions present in controls are shown in grey. The boxplot width is proportional to the number of deletions per group (see **Suppl. Figure S8**). P values were computed with independent 2-group Man-Whitney Unpaired Test.

**Suppl. Figure S10.**
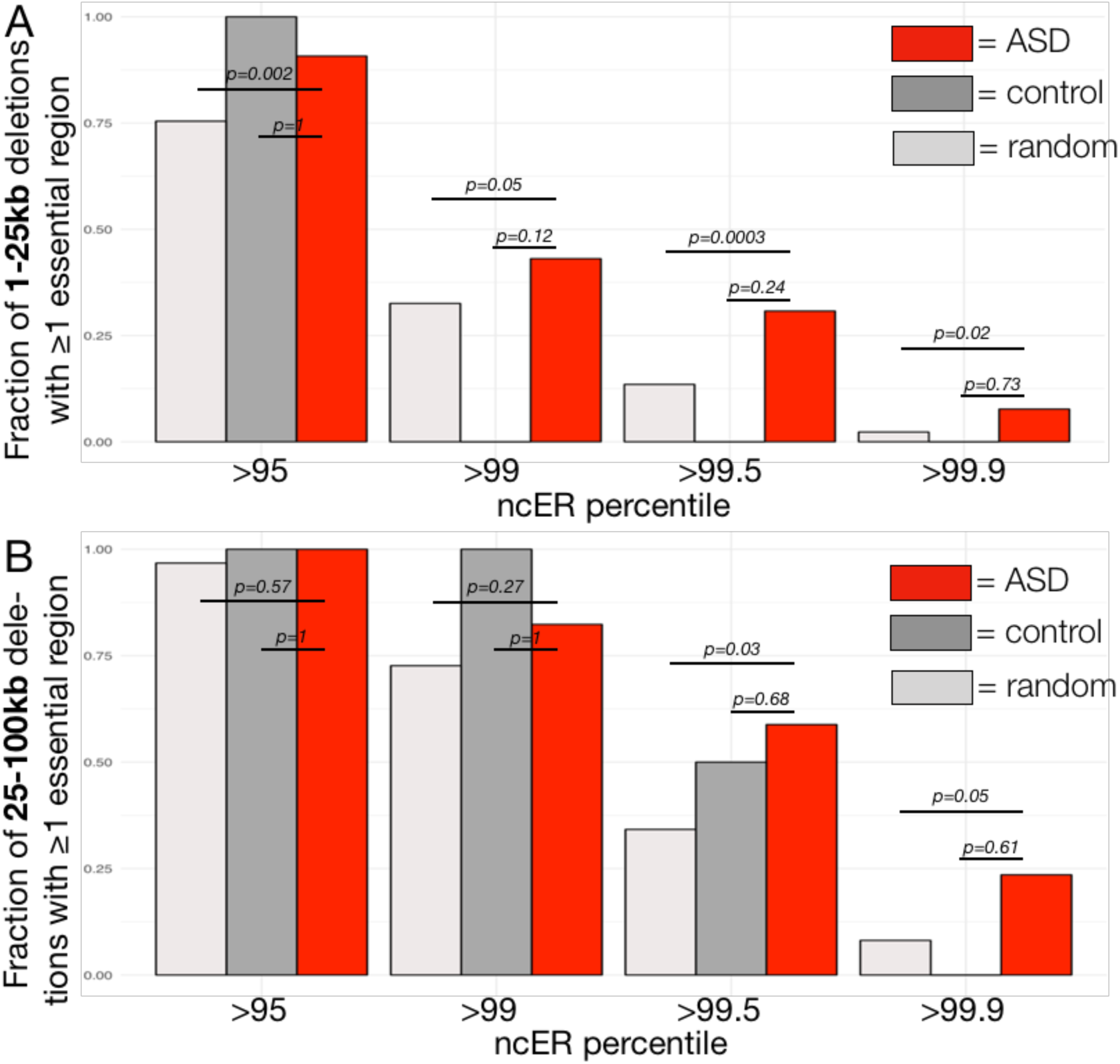
Fraction of transmitted deletions with essential functional domains. **Panel A.** Fraction of 1-25kb deletions with at least one essential bin. Essential bins are defined by four different ncER percentile thresholds. Autism and ASD deletions are shown in red (“ASD”, N=65), control deletions in dark grey (“control”, N=4) and random size-matched *in silico* deletions extracted genome-wide in light grey (“random”, N=6,900). **Panel B.** Fraction of 25-100kb deletions with at least one essential bin. Essential bins are defined by four different ncER percentile thresholds. Autism and ASD deletions are shown in red (“ASD”, N=17), control deletions in dark grey (“control”, N=2) and random size-matched *in silico* deletions extracted genome-wide in light grey (“random”, N=1,900).

**Suppl. Figure S11.**
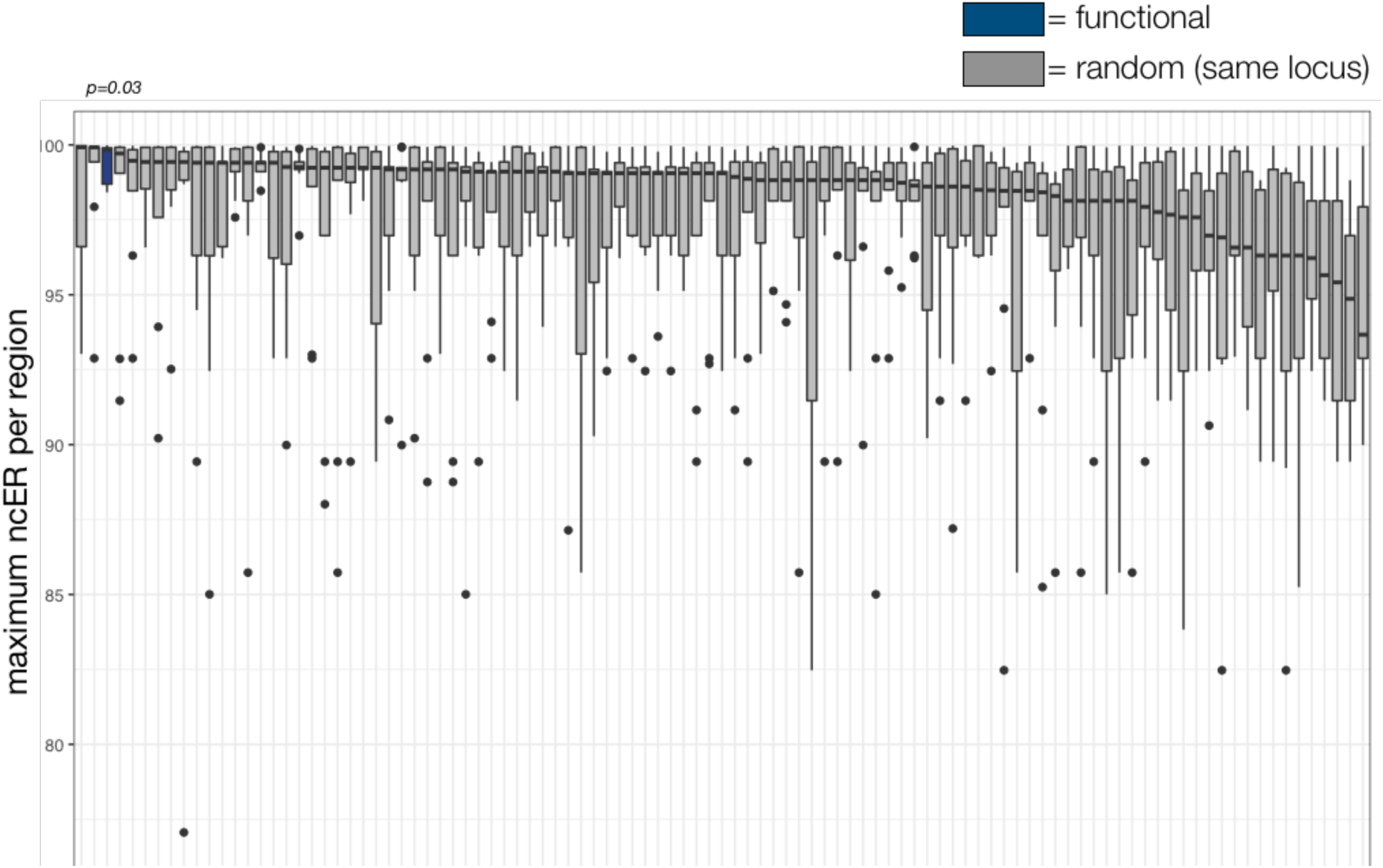
Enrichment in essential regions for mouse functional enhancers. Functional enhancers (N=9, blue) are placed among the highest ncER percentile distribution compared to 100 permutations (grey), each containing 9 regions matched by size to the enhancers and issued from the same genomic locus.

**Suppl. Table S1. Input feature description and accession links.**

Provided as separate file.

**Suppl. Table S2.**
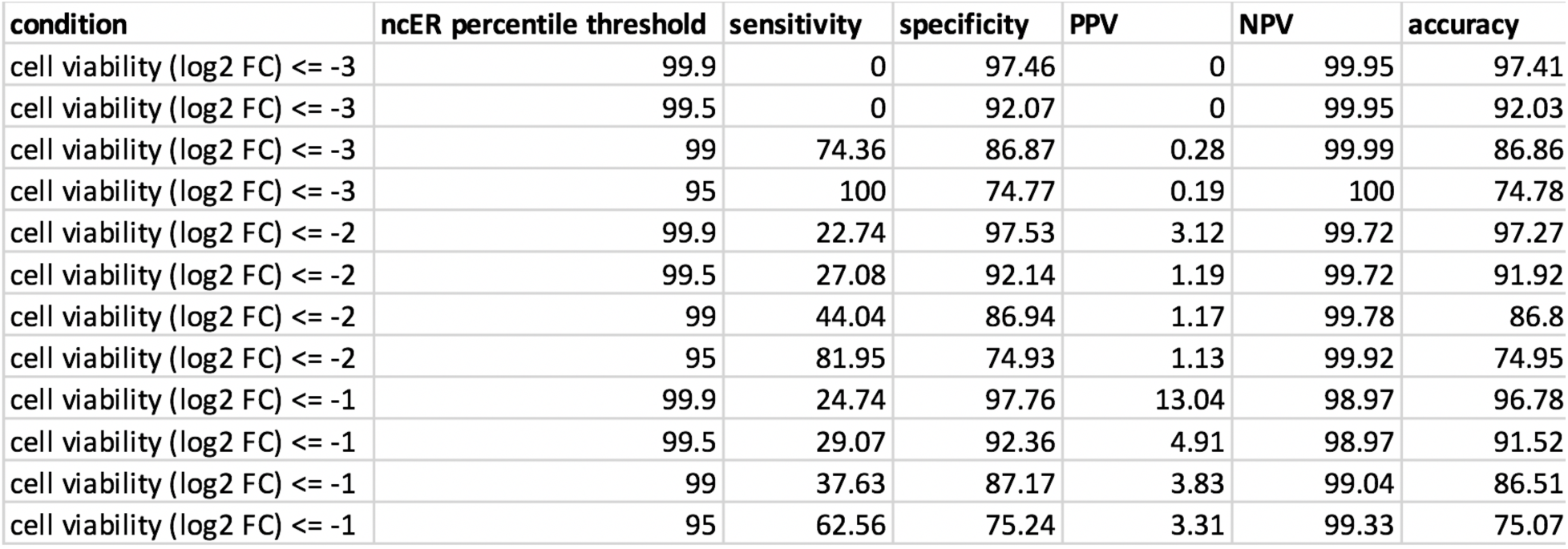
Predictive performance and accuracy of ncER compared to CRISPRi functional assays.

**Suppl. Table S3.** Predictive performance and accuracy of ncER compared to CREST-seq functional assays.

| condition | ncER percentile threshold | sensitivity | specificity | PPV | NPV | accuracy |
| --- | --- | --- | --- | --- | --- | --- |
| $-\log_{10}(\text{pval}) \geq 5.36$ | 99.9 | 0.00 | 96.07 | 0.00 | 99.52 | 95.63 |
| $-\log_{10}(\text{pval}) \geq 5.36$ | 99.5 | 26.42 | 76.46 | 0.51 | 99.56 | 76.23 |
| $-\log_{10}(\text{pval}) \geq 5.36$ | 99 | 50.94 | 64.03 | 0.65 | 99.65 | 63.97 |
| $-\log_{10}(\text{pval}) \geq 5.36$ | 95 | 92.45 | 19.02 | 0.52 | 99.82 | 19.36 |

